# A versatile dual-color bacterial reporter system highlights two distinct *Pseudomonas aeruginosa* Type 3 secretion system intracellular populations

**DOI:** 10.1101/2025.07.23.666013

**Authors:** Christopher J. Corcoran, David G. Glanville, Zachary J. Resko, Erin K. Cassin, Derrick L. Kamp, Karen L. Visick, Spencer V. Nyholm, Boo Shan Tseng, Abby R. Kroken, Andrew T. Ulijasz

## Abstract

Since their discovery, fluorescent reporters have revolutionized our ability to track gene and protein expression in real time. Ideally, two reporters are used, one constitutive signal for tracking viable bacteria and the other for measuring the expression of the gene/protein of interest. Unfortunately, these valuable tools are not available for most bacterial species, and if available are often not optimized for fluorophore protein folding rates and fluorescence intensity. Here we present a versatile dual reporter system, pCG-VmS, optimized for both transcriptional and translation fusions in Gram-negative bacteria. Using the important pathogen *Pseudomonas aeruginosa* for proof of concept, we demonstrate pCG-VmS utility in tracking transcriptional expression with flow cytometry and within a complex biofilm, and using a translational reporter fusion, monitor protein expression and visualize subcellular protein localization. We then analyzed T3SS-associated *exoS* toxin expression in infected host cells, which highlighted two distinct T3SS-dependent intracellular populations, one where the *exoS* promoter is turned on and the other where it is turned off in a smaller sub-population of bacterial cells (hereon referred to as T3SS-on and T3SS-off, respectively). Finally, we demonstrate the feasibility of spatiotemporal imaging in whole animals by using our system to monitor expression of an alternative sigma factor during *Vibrio fischeri* colonization of its squid host. Our findings demonstrate the versatile uses for the pCG-VmS vectors in microbiology, and that this vector can be used to visualize and separate distinct populations with precision for both *in vitro* and *in vivo* applications.

## Introduction

Fluorescent proteins (FPs) spanning the visible spectrum have been engineered to modify characteristics such as brightness, maturation time, excitation/emission spectra, and stability [1]. This has created a plethora of FPs to choose from, depending on the intended application and desired characteristics. In the field of molecular and cellular microbiology, FPs have shown great versatility and have been co-opted as cellular markers, as well as qualitative and quantitative experimental outputs [2]. This includes monitoring of gene expression [3, 4], determination of protein subcellular localization [5, 6], monitoring and tracking of distinct populations of microbial cells in complex mixtures [7, 8], assessing protein-protein interactions [9, 10], monitoring metabolite concentrations [11, 12], and tracking gene expression spatiotemporally during infection [13]. Due to their diversity and extensive applications, the use of FPs has become integral to studying and interrogating cellular dynamics and functions.

Measurement of gene expression at the transcriptional level is a widely employed approach to assess the response of microbial cells to environmental conditions or genetic manipulations [14, 15]. Changes in gene expression can be assessed directly at the level of mRNA through Northern blots, RT-qPCR, or RNA-sequencing [16]. However, these techniques require the isolation of often unstable mRNA from microbial cells, as well as multiple downstream steps before the data can be analyzed. Further, monitoring gene expression during microbial infection using these aforementioned techniques can be even more challenging and requires sacrificing animals at the desired timepoints. Considering these obstacles, genetically encoded transcriptional or translational reporters are often preferentially used in microbial systems to assess promoter activity or protein expression levels and cellular localization of specific proteins. When using a single reporter system, the variable signal must be normalized to the number of viable cells being measured, which is most often done spectrophotometrically through an optical density measurement or direct enumeration of colony forming units. These readings can become unreliable as cells can clump and/or may form biofilms, causing erratic changes in the normalization value that then affects the final value calculated for the gene expression. One way to solve this problem is to use a second, constitutively expressed FP with an excitation and emission that does not spectrally overlap the variable FP (experimental) output. This second signal can then be used to normalize the experimental fluorophore signal, allowing for more accurate quantitation of expression [17]. Furthermore, for plasmid-based systems, the constitutively expressed FP also allows for normalization to plasmid copy number, which can add additional variance between samples [18, 19].

FPs are also often translationally fused to a gene of interest to determine the precise subcellular localization of a protein. While most bacteria do not contain membrane bound organelles, it has become increasingly apparent that subcellular organization of proteins and protein-complexes occurs in bacteria more than what was previously thought [20–22]. In order to precisely determine the subcellular localization of a protein, the exact boundary and shape of the bacterium must also be visualized. This can be accomplished by fluorescent-membrane staining [23], phase contrast (PC) [24, 25], differential interference contrast (DIC) [26], or expression of an evenly-distributed cytoplasmic fluorescent protein [27]. One advantage of using a cytoplasmic fluorescent protein is that additional cellular manipulations such as membrane staining or additional microscopy technology (e.g. PC) are not necessary to obtain the shape of the bacteria before imaging.

While the dual fluorescent reporter offers a superior option for *in vitro* assays, for *in vivo* infections they are considered the ‘holy grail’ of imaging tools. However, several obstacles exist for fluorescent dual reporters for use with whole-animal, spatiotemporal imaging. Ideally when possible, an optically-accessible animal model system is preferred, such as zebrafish [13]. For less optically accessible animals, such as mice, special care must be taken in selecting FPs with longer excitation and emission wavelengths (ideally near infrared) that are able to traverse dense tissue without significant absorption [28, 29].

When building fluorescent reporter systems, one must consider many factors, including promoter strength, and stability and toxicity of the FP in question. For example, the use of dual fluorophore vectors in the highly studied opportunistic pathogen *Pseudomonas aeruginosa* often leads to growth defects and toxic effects [30]. Thus, to overcome these obstacles, a dual reporter system requires comparably strong signals from both the constitutively expressed “normalization” FP signal and the tightly controlled (experimental) FP signal which is expected to show measurable variability. Further, multiple FPs to be measured simultaneously must be optimized for relatively comparable stability and resistance to photobleaching to enable longitudinal imaging studies in both cell culture and whole-animal imaging. Most importantly, one must consider two FPs that have minimal spectral overlap and have standard excitation and emission wavelengths that can be detected using most plate readers, flow cytometry systems, various microscopy techniques, and *in vivo* whole animal imaging systems.

Herein, we describe two broad-host-range expression cassettes for gram-negative bacteria (the pCG-VmS system), including *P. aeruginosa*, that can be easily transferred into existing vector-based systems to enable both transcriptional and translational fusions. We verified the utility of these vectors by using transcriptional and translational gene fusions for the *P. aeruginosa* aldehyde-responsive gene *arqI*, which is tightly and specifically controlled by the presence of the aldehyde glyoxal (GO), which drives ArqI polar localization upon GO exposure [31]. In addition, we utilized this system for the first time to visualize, track, and quantify the formation of subpopulations of Type 3 Secretion System (T3SS) positive (T3SS-on) and negative (T3SS-off) bacteria during intracellular infection of corneal epithelial cell monolayers [32]. Finally, we tested the use of our dual reporter system with whole animal imaging and visualized gene expression of an alternative sigma factor during *Vibrio fischeri* colonization of *Euprymna scolopes* Hawaiian bobtail squid light organ. Taken together, we anticipate the pCG-VmS dual reporter vectors will enhance the overall precision and accuracy of basic reporter data collection as well as longitudinal imaging techniques when studying gene and protein expression gram-negative bacteria.

## Results

### Design and characterization of pCG-VmS

To develop a plasmid system that would have wide utility in monitoring expression, at either the transcriptional or translational level, we utilized the plasmid pCC21 as a backbone structure [33]. pCC21 maintains a low copy number (approximately 5-10 copies per cell) through the pBBR1 origin [34]. For reporter gene output after an initial FP screening process, the engineered monomeric red fluorescent protein (RFP) mScarlet-I was selected due to its inherent high brightness and low background fluorescence (resulting from its red-shifted emission wavelength), which allows for increased sensitivity when compared to other more blue-shifted FPs [35]. The mScarlet-I variant of mScarlet also possesses a more rapid maturation time when compared to other RFP variants [36], making it a great candidate for a reporter gene that can maintain sensitivity while enabling rapid detection. Superfolder GFP (sfGFP) was then selected as a constitutive fluorescent signal due to its similar folding time and enhanced brightness [36]. To constitutively express sfGFP at levels amenable to most detection needs, a synthetic *lac*-based promoter, P_A1/04/03_ [37, 38], was originally chosen for its ability to drive high levels of expression, and its previous use as a constitutively expressed promoter for fluorophores in *P. aeruginosa* [38–40]. However, preliminary testing with this promoter driving sfGFP expression resulted in a disruptive bacterial cell filamentation phenotype – a phenomenon that occurred when mScarlet-I was simultaneously expressed at high level using the promoter for *exoS* (**Movie S1**). Alternatively, a subpopulation of bacteria was observed that suppressed expression of sfGFP (seen within the same field containing filamentous bacteria in **Movie S1**). Indeed, it has been previously observed that in *P. aeruginosa*, high expression of two fluorophores can lead to unwanted growth or virulence defects, or mutagenesis and/or loss of the vector to suppress expression of one or both fluorophores [41]. To circumvent this problem, the P_A1/04/03_ promoter was exchanged for the less active *lacUV5* promoter, which also exerts its expression independent of CRP control [37]. After this alteration, the sfGFPintensity was reduced, alleviating the cell filamentation phenotype that occurred with simultaneous mScarlet-I expressed from a strong promoter (**Movie S1**). The finalized vector was named pCG-VmS (<u>C</u>onstitutive <u>G</u>FP, <u>V</u>ariable <u>mS</u>carlet-I) (**Fig 1A**).

**Figure 1.**
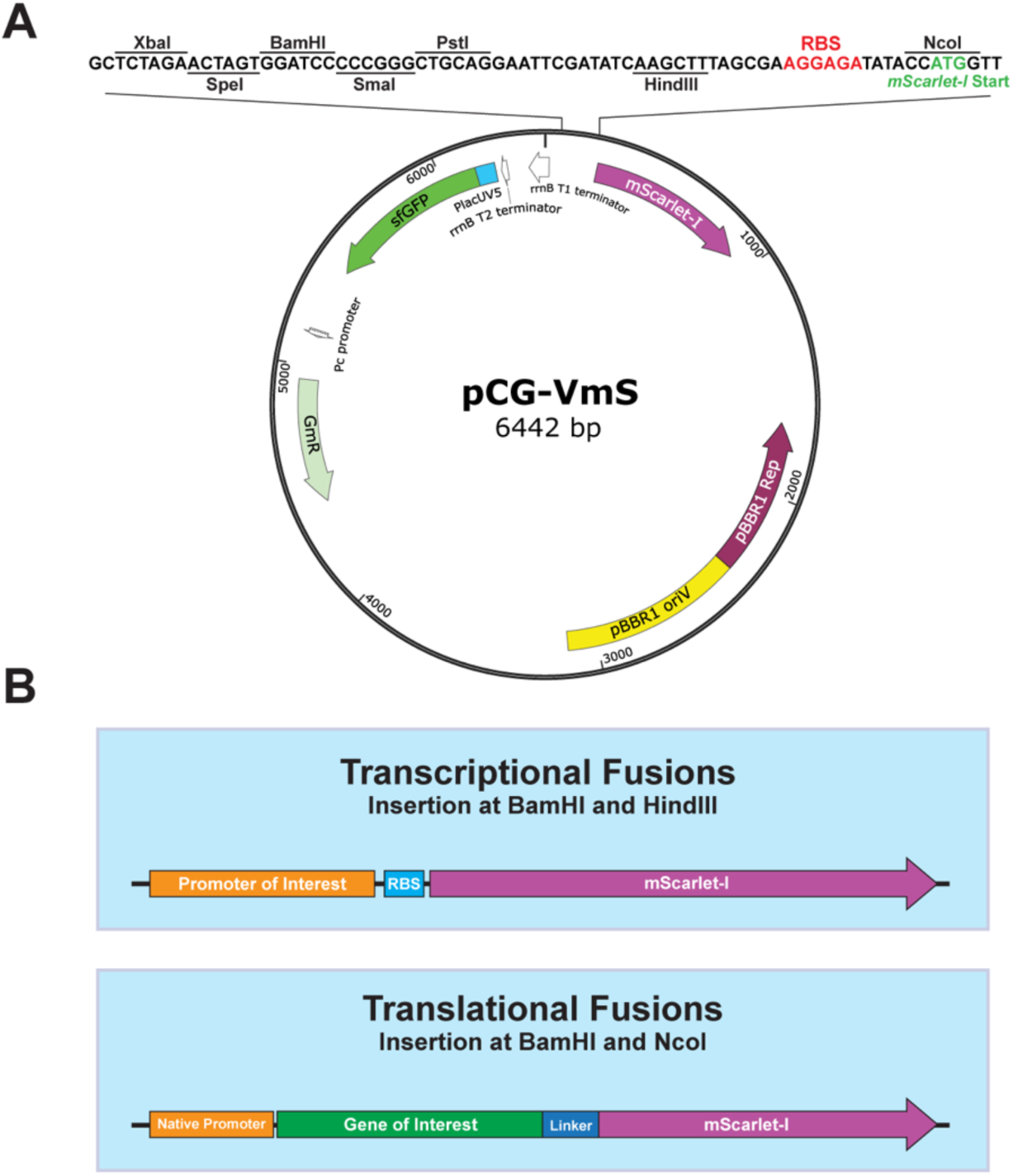
pCG-VmS design. (**A**) MCS and plasmid map of pCG-VmS. The ribosome binding site (RBS) and mScarlet-I start codon are highlighted in red and green, respectively. The plasmid map was generated using SnapGene. (**B**) Schematic of transcriptional and translational fusions that can be made depending on which restriction enzymes are utilized for insertion.

The multiple cloning site (MCS) of pCG-VmS was designed for easy insertion of promoters upstream of mScarlet-I (with its associated RBS) to generate transcriptional fusions (**Fig. 1A, B**). For translational fusions, the RBS can be removed by digestion with *Nco*I and another selected upstream restriction enzyme site. The gene of interest and its upstream regulatory region can then be inserted in frame with *mScarlet-I* with a 3’ linker region - since the *Nco*I site contains the mScarlet-I start codon to create a C-terminal mScarlet-I fusion protein (**Fig. 1B**). In addition, Rho-independent *rrnB* T1 and T2 terminators [42] were inserted upstream of the MCS to prevent transcriptional readthrough into the *mScarlet-I* promoter.

### pCG-VmS assessment during *in vitro* growth

Relative to its non-fluorescent parent plasmid pCC21, pCG-VmS caused no deleterious effects on growth of WT *P. aeruginosa* MPAO1 (**Fig. 2A**). As stated before, the low background fluorescence at red-shifted wavelengths allows for increased sensitivity of RFPs, such as mScarlet-I [43]. However, cells harboring pCG-VmS did exhibit some detectable mScarlet-I fluorescence throughout 8 hours of growth compared to the background levels of the non-fluorescent control; **Fig. 2B**), indicating, as would be expected with any reporter system, a basal level of mScarlet-I signal is indeed produced despite the introduction of the terminators. Robust sfGFP signal well above the background intrinsic fluorescence of the cell and complex media was also seen (**Fig 2C**; ∼100-fold increase in sfGFP channel fluorescence after 8 hours when compared to the non-fluorescent control). In addition, the raw fluorescence values of the sfGFP signal closely mimicked the growth of the cells when measured by the absorbance at 600 nm (**Figs. 2A, C**). When the sfGFP signal was normalized to the OD_600_ value, a noticeable decrease in the fluorescence density was seen as the cells entered the logarithmic growth phase (**Fig. 2D**). One likely explanation for this phenomenon could be dilution of the sfGFP molecules as the cells divide rapidly [44], and a slight delay in fluorescence of newly synthesized sfGFP that must first mature. Similarly, a decrease in fluorescence density was also seen when monitoring the mScarlet-I signal (**Fig. 2D**). Since both FPs should theoretically be equally affected by this decrease, normalization of the mScarlet-I signal to the sfGFP signal would therefore be a preferred form of normalization rather than the OD; a superiority that is supported by the literature [8].

**Figure 2.**
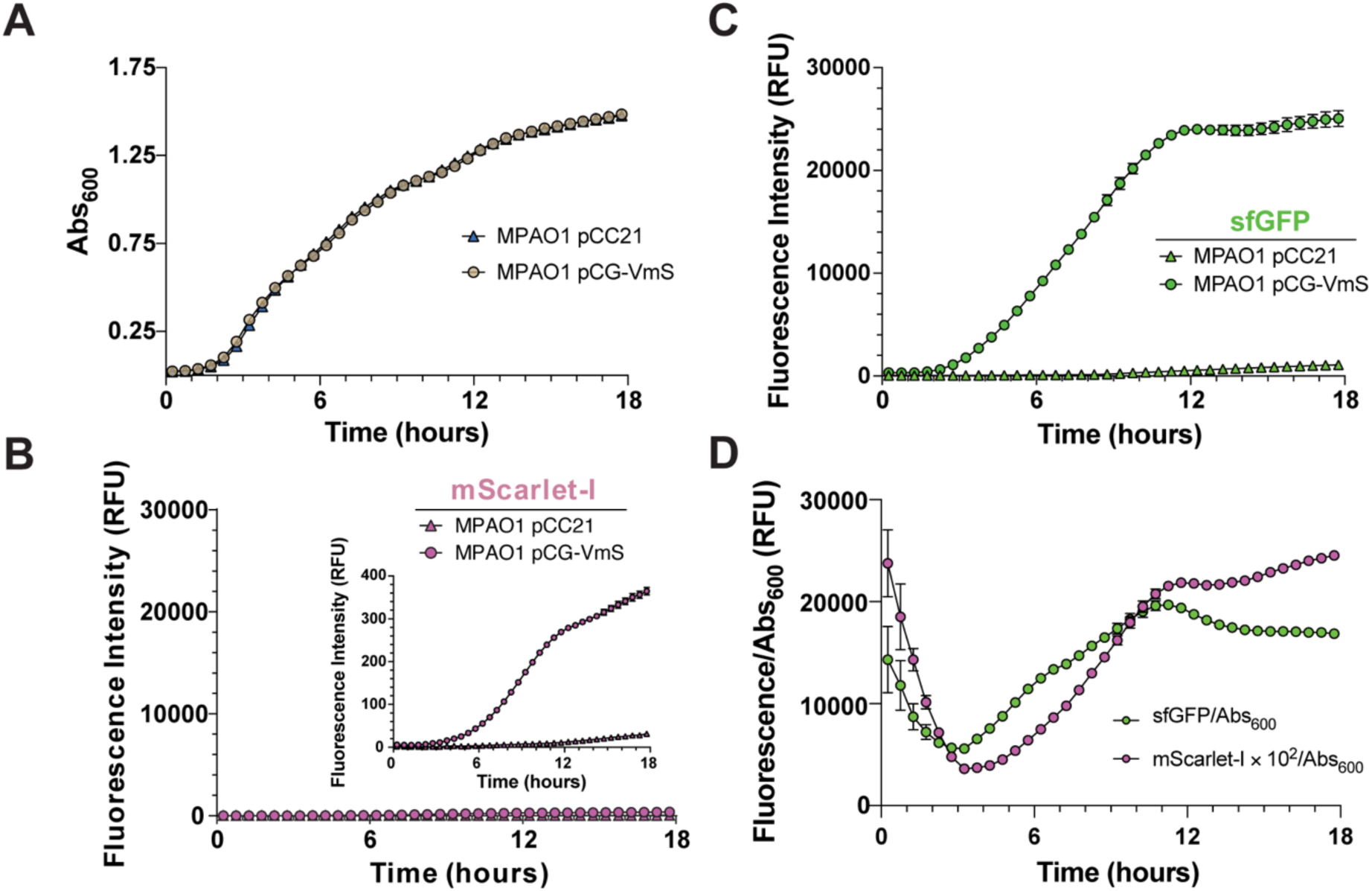
Characterization of pCG-VmS during planktonic growth. *P. aeruginosa* MPAO1 harboring pCG-VmS or the non-fluorescent control plasmid pCC21 was grown in LB medium with antibiotics for 18 hours. (**A**) Absorbance at 600 nm, (**B**) raw mScarlet-I signal, and (**C**) raw sfGFP fluorescence signal over time is shown. (**D**) mScarlet-I and sfGFP signals were normalized using the absorbance at 600 nm. Normalization of mScarlet-I and sfGFP signals to Abs600 results in a similar “S”-shaped curve that could be due to fluorescence dilution and/or fluorophore maturation.

### Utility as a transcriptional reporter in planktonic and biofilm forming *P. aeruginosa*

We recently demonstrated that *arqI-gloA2* operon transcription is tightly controlled and specifically induced upon exposure of *P. aeruginosa* to the toxic aldehyde GO, but not by other aldehydes such as methylglyoxal (MGO) [31, 33] (**Fig. 3A**). Due to this tight control and inherent specificity, we exploited the *arqI* promoter to test the ability of pCG-VmS to perform as a transcriptional reporter. 145 base pairs of DNA sequence upstream of the *arqI* start codon was cloned into pCG-VmS to create a transcriptional reporter for the *arqI-gloA2* operon to yield the plasmid pCG-P*_arqI_*-mS. In line with our previous work [33], we found that transcription from this promoter was highly controlled and specific to GO. Increasing concentrations of GO caused a dose-dependent increase in reporter activity, an effect that did not occur upon MGO addition until the highest concentration tested (**Fig. 3B**). pCG-P*_arqI_*-mS showed a high level of induction, with an approximate 100-fold change in normalized reporter activity at 4 mM GO when compared to untreated control. Notably, our previous studies utilizing P*_arqI_* transcriptional fusions to a fluorescent reporter found only a 10-fold increase in promoter activity upon treatment of 4 mM GO [33], suggesting that when normalized to sfGFP signal, the mScarlet-I signal from the dual pCG-P*_arqI_*-mS system may exhibit increased sensitivity due to lower background interference when compared to previous, single reporter constructs [31]. After GO addition, reporter activity increased rapidly, with detectable levels of mScarlet-I fluorescence above baseline, appearing 30 minutes post-GO addition and increasing over a period of 3.5 hours (**Fig, 3C**, left). Of note, insertion of the *arqI* promoter reduced the mScarlet-I signal below the level of the empty vector control, suggesting repression of the promoter by an unknown transcription factor (**Fig. 3C**, left). Important for pCG-VmS utility, we could not detect any growth defects even after the highest levels of mScarlet-I signal were achieved, indicating negligible cytotoxicity (**Fig. 3C**, right).

**Figure 3.**
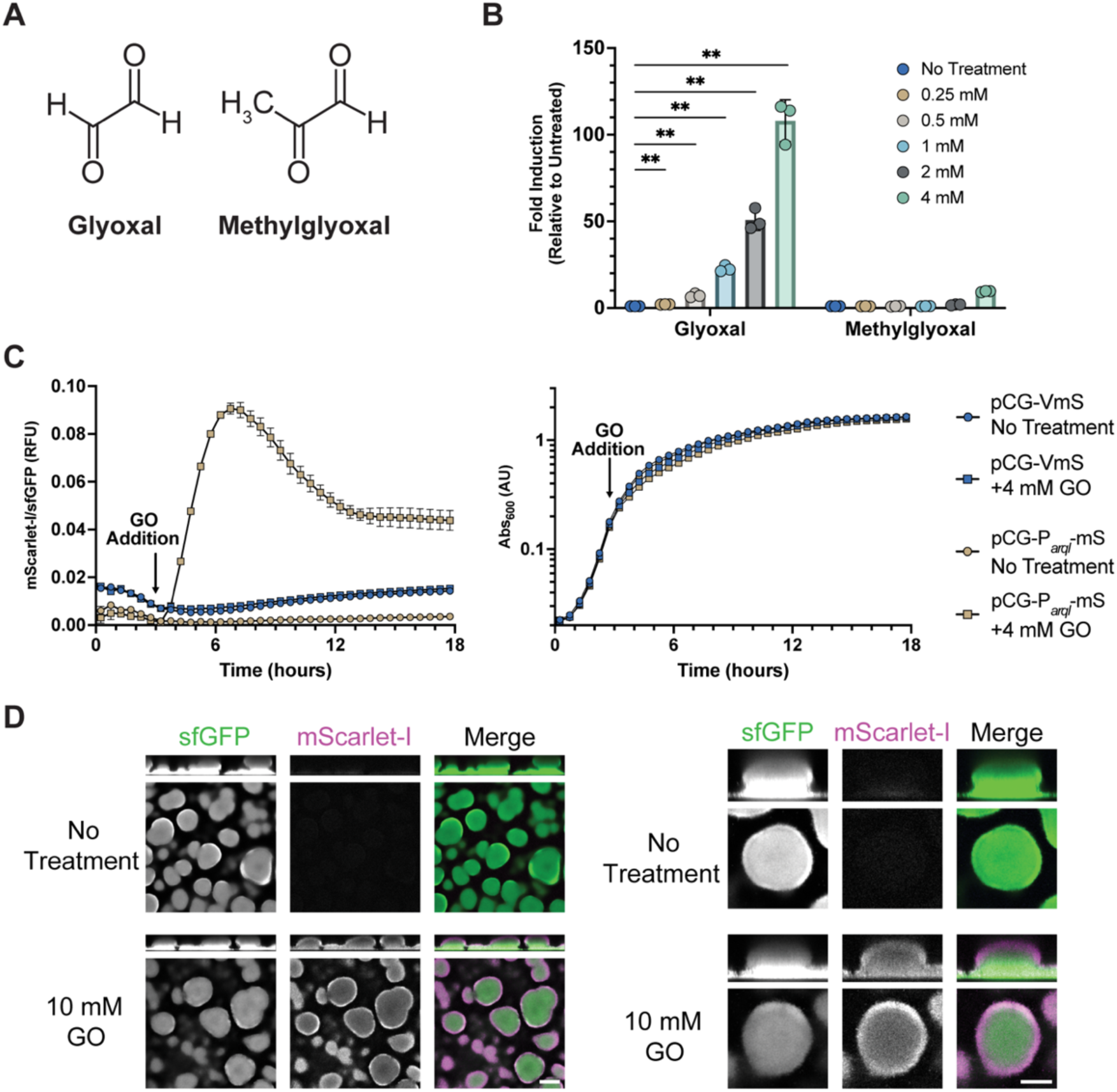
pCG-VmS transcriptional reporter utility. (**A**) Chemical structures of GO and MGO. (**B**) Dose response of pCG-P*arqI*-mS to GO and MGO at three hours post addition. Values are expressed as mScarlet-I/sfGFP relative to the no treatment control. *, P < 0.05; **, P < 0.01; ***, P < 0.001 by a one-sample *t* test. (**C**) Normalized mScarlet-I fluorescence signal (left) and growth according to optical density reading (right) of pCG-P*arqI*-mS after 4 mM GO was added at 3 hours post *P. aeruginosa* inoculation. (**D**) Induction of the *arqI* promoter by 10 mM GO treatment in biofilms grown under flow. Scale bar = 100 μm (left) or 50 μm (right).

### Use of pCG-VmS to observe promoter expression in biofilms using a flow cell

We next sought to investigate the use of this reporter system in more complex three-dimensional environments by exploiting flow cell technology. WT MPAO1 biofilms were grown in a flow cell in 1% TSB media for 96 hours before the addition of 10 mM GO. After 3 hours of GO incubation, the mScarlet-I signal increased, with a noticeable heavier induction around the periphery of the biofilm (**Fig. 3D**). This suggests GO may have a limited ability to penetrate the biofilm mass, albeit not completely, and confirms the utility of this reporter plasmid for imaging biofilms.

### Use of pCG-VmS to visualize promoter expression at the single cell level with flow cytometry

To visualize the dynamics of *arqI-gloA2* promoter induction at the single cell level, flow cytometry was used. Relative to the non-fluorescent parent plasmid pCC21, the pCG-P*_arqI_*-mS harboring strain exhibited robust sfGFP fluorescence in roughly 99.5% of cells analyzed by flow cytometry (**Fig. 4A**). Conversely, in the absence of GO, minimal levels of mScarlet-I signal were seen, indicating some expected basal level of *arqI* promoter activity under standard culture conditions (**Fig. 4B**). When increasing concentrations of GO were added, the *arqI-gloA2* promoter showed a graded response (**Fig. 4A, B**). The addition of 0.5 mM GO resulted in a wide distribution of mScarlet-I signal, suggesting differential induction occurring across the population of bacteria (**Fig. 4B**). As the concentration of GO increased, the population exhibited increasingly homogenous expression of mScarlet-I as indicated by a sharper peak, culminating in 95% induction at 2 mM and 99% induction at 4 mM GO (**Fig. 4B**).

**Figure 4.**
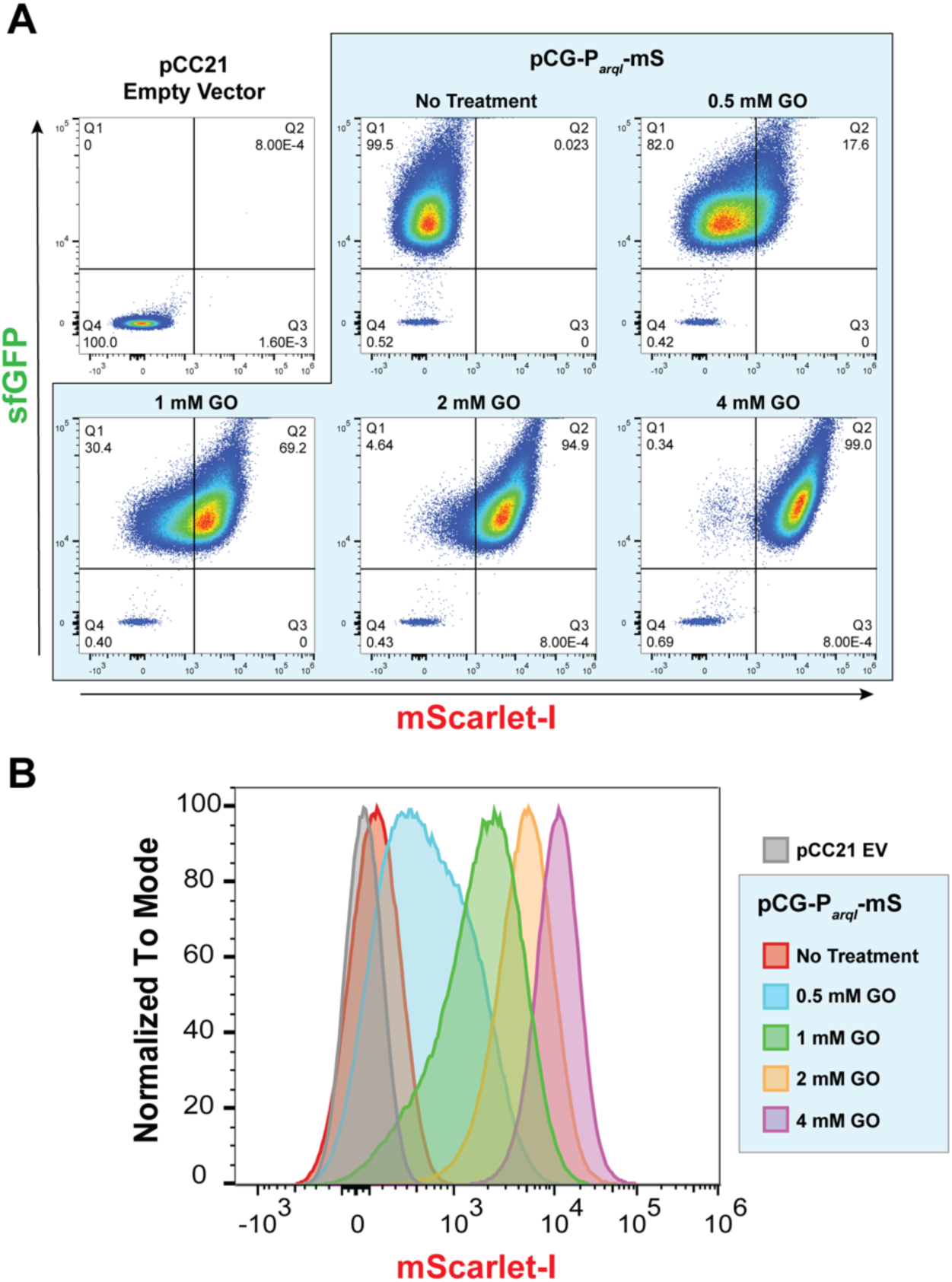
Population dynamics of *arqI* expression in response to GO. (**A**) Pseudo-colored dot plots of mScarlet-l expression upon increasing concentrations of GO added to MPAO1 pCG-P*arqI*-mS cells depicting sfGFP-positive population and mScarlet-l induction by GO. (**B**) Flow cytometry histogram of data presented in *A*. Populations are normalized to mode. One representative of three biological replicates is shown. MPAO1 pCC21 was used as a non-fluorescent control. EV, empty vector.

### pCG-VmS utility as a translational reporter for protein cellular localization

To test the utility of pCG-VmS for use with translational fusions and subcellular protein localization, the *arqI* promoter and coding sequences were inserted in-frame with *mScarlet-I* with a linker to form a C-terminal fusion protein [27]. WT MPAO1 cells harboring this fusion vector were spotted onto 1.5% agarose LB pads and imaged over the course of 3.5 hours. In the absence of GO, no localization or robust fluorescent signal was seen, only the constitutive sfGFP signal that outlined the shape of dividing bacteria (**Fig. 5**, above). However, as we previously observed with a sfGFP fusion to ArqI [31], upon GO induction, the ArqI-mScarlet-I fusion protein clearly localized to the cellular pole in a subpopulation of bacteria (**Fig. 5**, below). These results show that pCG-VmS can also be used to detect bacterial cell protein localization using translational fusions.

**Figure 5.**
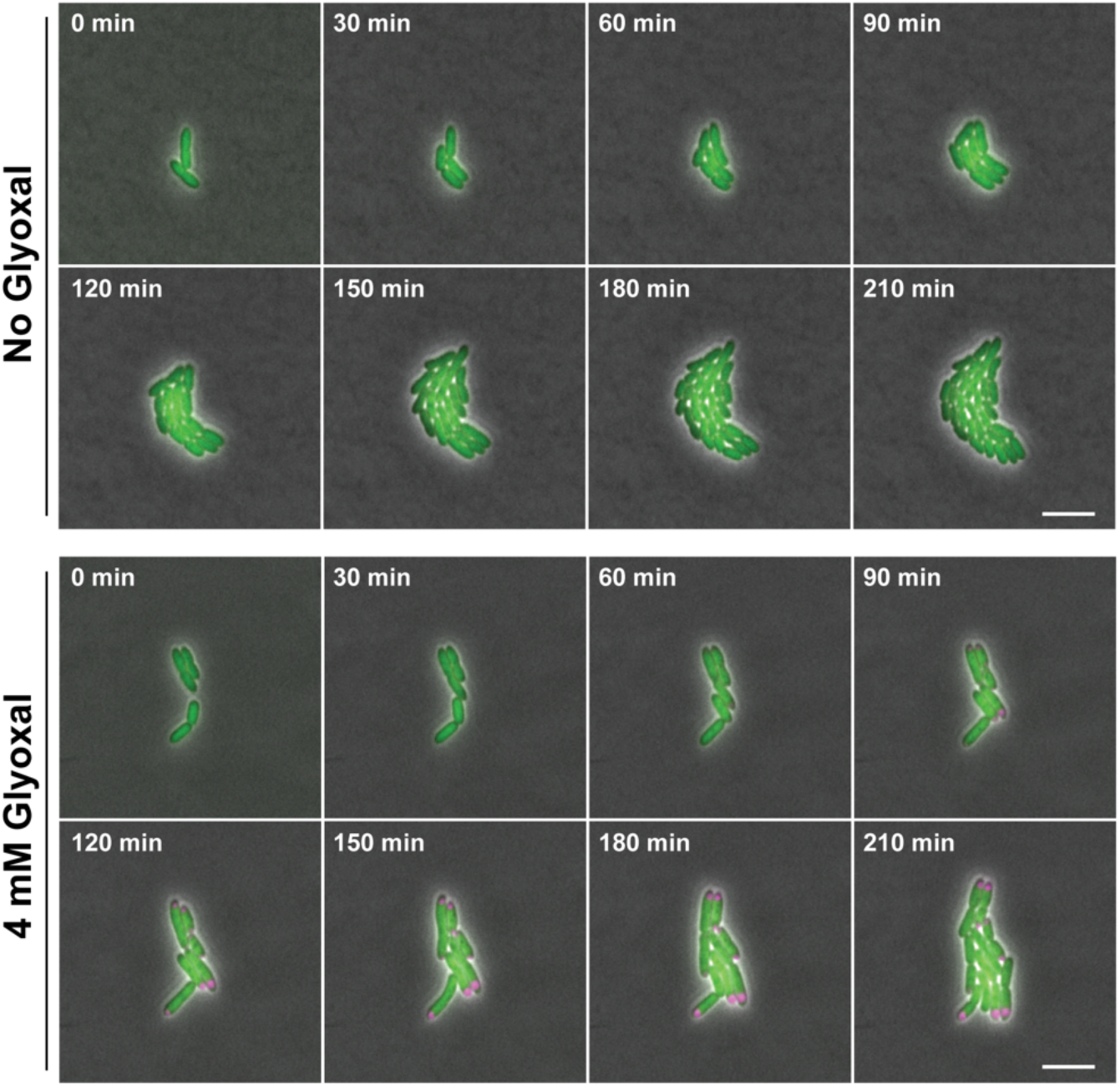
Polar localization of ArqI following induction by GO. Time-lapse microscopy of MPAO1 pCG-P*arqI*-ArqI-mS cells cultured to mid-logarithmic phase before transfer to an 1.5% agarose pad without (above) or with 4 mM GO (below) for imaging. Scale bar = 5 μm.

### Visualization of T3SS positive (on) and negative (off) populations during intracellular infection

*P. aeruginosa* has the ability to invade epithelial cells [45] and establish a cytoplasmic niche [41, 46].

After initial internalization, *P. aeruginosa* can activate the T3SS to escape the vacuole [41] and freely replicate inside the cytoplasm until subsequent host cell death [47]. Alternatively, a subpopulation of vacuolar *P. aeruginosa* has been observed to remain T3SS-off, persisting inside vacuole compartments [48]. However, genetic tools have not been available to visualize these two distinct populations simultaneously and give conclusive evidence of these suspected virulent and quiescent subpopulations. Moreover, these T3SS-on and -off populations could not be accurately quantified. One tool used for the T3SS-on cytoplasmic population, pJNE05, is a GFP-transcriptional reporter for the T3SS toxin ExoS that acts as an output indicator for T3SS activation [49], but does not allow for visualization of the T3SS-off bacterial population during infection. While arabinose-inducible GFP vectors have also been explored for these purposes, they were found to suppress T3SS activity [50] and have other general limitations in *P. aeruginosa* [51]. To help with these issues, we constructed a transcriptional reporter in pCG-VmS to the *exoS* promoter, pCG-P*_exoS_*-mS.

In comparison to pJNE05, pCG-P*_exoS_*-mS showed similar reporter induction and kinetics when T3SS was induced using the calcium chelation agent EGTA, as measured by the mScarlet-I/sfGFP ratio over time (**Fig. 6A**). As expected, we observed a lower background signal with the pCG-P*_exoS_*-mS data, likely due the more accurate method of utilizing a second FP signal (sfGFP here) to normalize. In early log phase (lag phase to approximately OD_600_ 0.3), a slight delay in reporter activity was seen with pCG-P*_exoS_*-mS, which could be due to the marginally longer maturation time of mScarlet-I when compared to GFP [36]. To assess the utility of pCG-P*_exoS_*-mS during infection, we exposed telomerase-immortalized human corneal epithelial cells (hTCEpi) to WT PAO1 harboring either pJNE05 or pCG-P*_exoS_*-mS. T3SS activation could be visualized within 4.5 hours post infection, with expansion of the T3SS-positive population being comparable between the two strains (**Fig. 6B**, **Movie S2**). However, use of our pCG-P*_exoS_*-mS plasmid also allowed for visualization of the expanding T3SS-off populations when extracellular fluorescent bacteria are cleared with the addition of polymyxin B (**Fig. 6C**, **Movie S3**). Some T3SS-off bacteria resided inside the same cells as expanding T3SS-on bacteria, as previously observed [48]. Detection of this previously untraceable T3SS-off population using pCG-P*_exoS_*-mS verifies evidence of their existence from prior studies [52] and therefore now pave the way for mechanistic investigations of processes dictating vacuolar exit to be performed. Together, these data validate the use of pCG-VmS for tracking discrete bacterial populations in the context of host cell infections.

**Figure 6.**
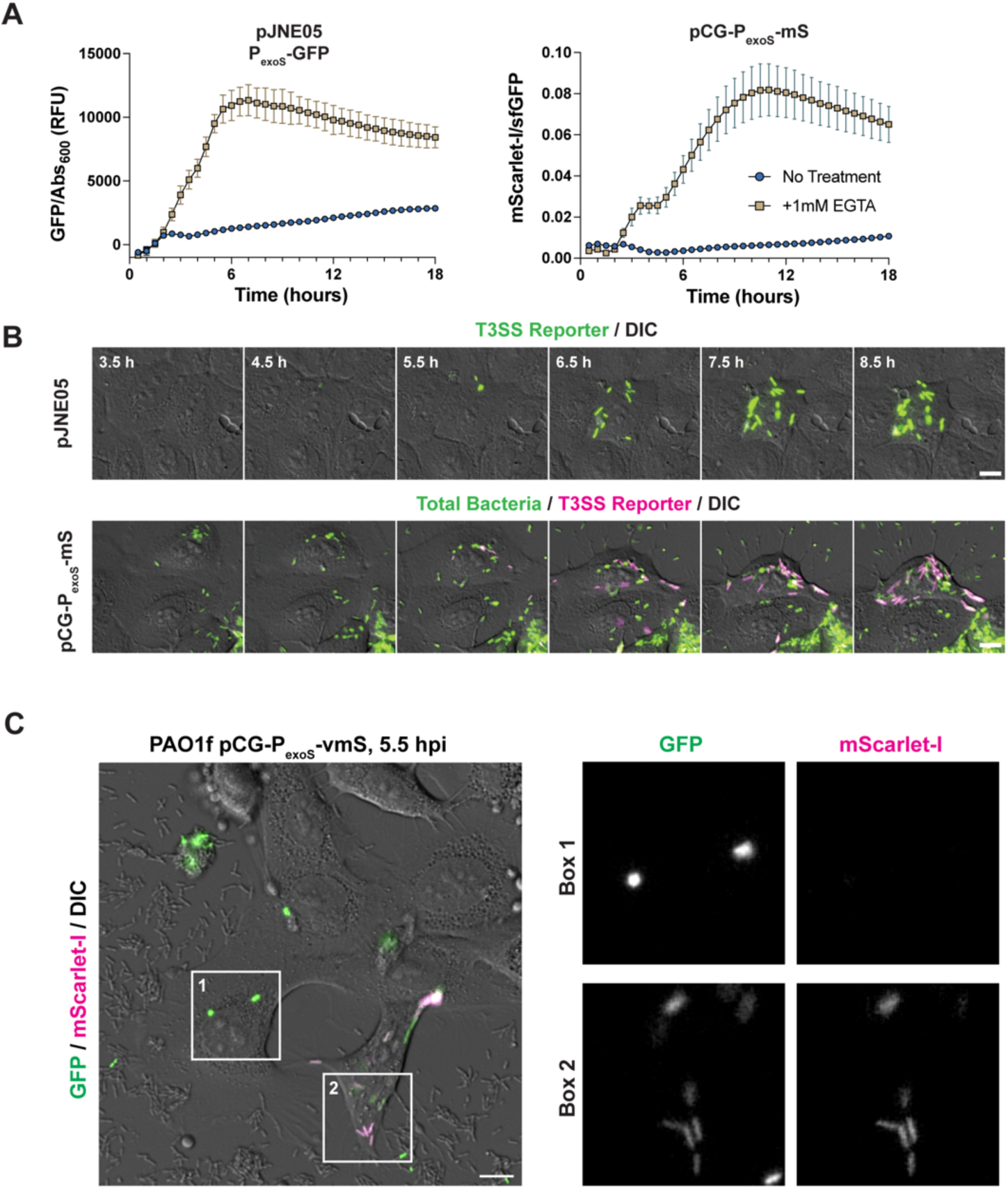
Induction of pCG-PexoS-mS by EGTA and Infection of Corneal Epithelial Cells. (**A**) Normalized fluorescent signals of EGTA-mediated induction of the *exoS* promoter in PAO1f utilizing pJNE05 (left) and pCG-P*exoS*-mS (right). (**B**) *In vitro* infection of hTCEpi corneal epithelial cells comparing the kinetics of T3SS activation between pJNE05 (above) and pCG-P*exoS*-mS (below) in PAO1f. Scale bar = 10 μm. (**C**) *In vitro* infection of hTCEpi corneal epithelial cells invaded by PAO1f with pCG-P*exoS*-mS shown at 5.5 hours post infection. At 3 hours post infection, extracellular bacteria were killed with amikacin and polymyxin B, which diminishes constitutive sfGFP fluorescence of extracellular bacteria by 5 hours post infection. Populations of intracellular T3SS-off (box 1) and T3SS-on (box 2) bacteria are shown.

### *In vivo* imaging of *V. fischeri*-colonized juvenile *E. scolopes* squid

Due to the low background fluorescence and high brightness of the mScarlet-I reporter, we hypothesized that the pCG-VmS reporter cassette would allow for robust *in vivo* imaging in an optically accessible animal model. One animal with a relatively optically clear body is the Hawaiian bobtail squid, *Euprymna scolopes*, whose symbiotic light organ is colonized by the bacterium *V. fischeri* [53, 54]. As a result of the light organ properties, it is possible to achieve spatiotemporal visualization of *Vibrio fischeri* reporter constructs as the bacteria colonizes the light organ. To capture these dynamics, dual fluorescence vectors have been utilized for *in vivo* imaging of colonized juvenile *E. scolopes* squid before [55, 56]. To determine if our dual reporter system would work in *V. fischeri*, we transferred the fluorescence cassette from pCG-VmS into the *Vibrio* specific plasmid, pVSV105, creating pVSV105-CG-VmS. Due to differences in sfGFP expression in *V. fischeri*, we utilized the constitutive P_A1/04/03_ promoter [12] in the pVSV105-CG-VmS vector to increase the sfGFP signal to the required more robust levels.

After creating pVSV105-CG-VmS, we sought to first verify its utility in *V. fischeri in vitro*. We constructed a transcriptional reporter using the promoter of *cysK* (pVSV105-CG-P*_cysK_*-mS), which is highly repressed in *V. fischeri* in the presence of cystine [57]. When grown in minimal media supplemented with cystine, an ∼10-fold decrease in signal was detected compared to media without any cystine supplementation (**Fig. 7A**). This decrease in reporter activity is in agreement with previous results of *cysK* promoter activity [57], and confirmed the utility of pVSV105-CG-VmS as a viable *in vitro* transcriptional reporter. We then tested pVSV105-CG-VmS as a reporter system for whole animal imaging. For these experiments, we used the promoter for the alternative sigma factor *rpoQ* (plasmid pVSV105-CG-P*_rpoQ_*-mS) for two reasons: (i) *rpoQ* encodes an unusual sigma factor whose overexpression impacts motility, luminescence, and chitinase activity (phenotypes that are relevant to the interaction of *Vibrio fischeri* with its symbiotic host [58]), and [ii] transcription from the *rpoQ* promoter is potentially controlled by the LitR transcription factor [59] (information that provides a starting point for future evaluation of the importance of specific promoter sequences).

**Figure 7.**
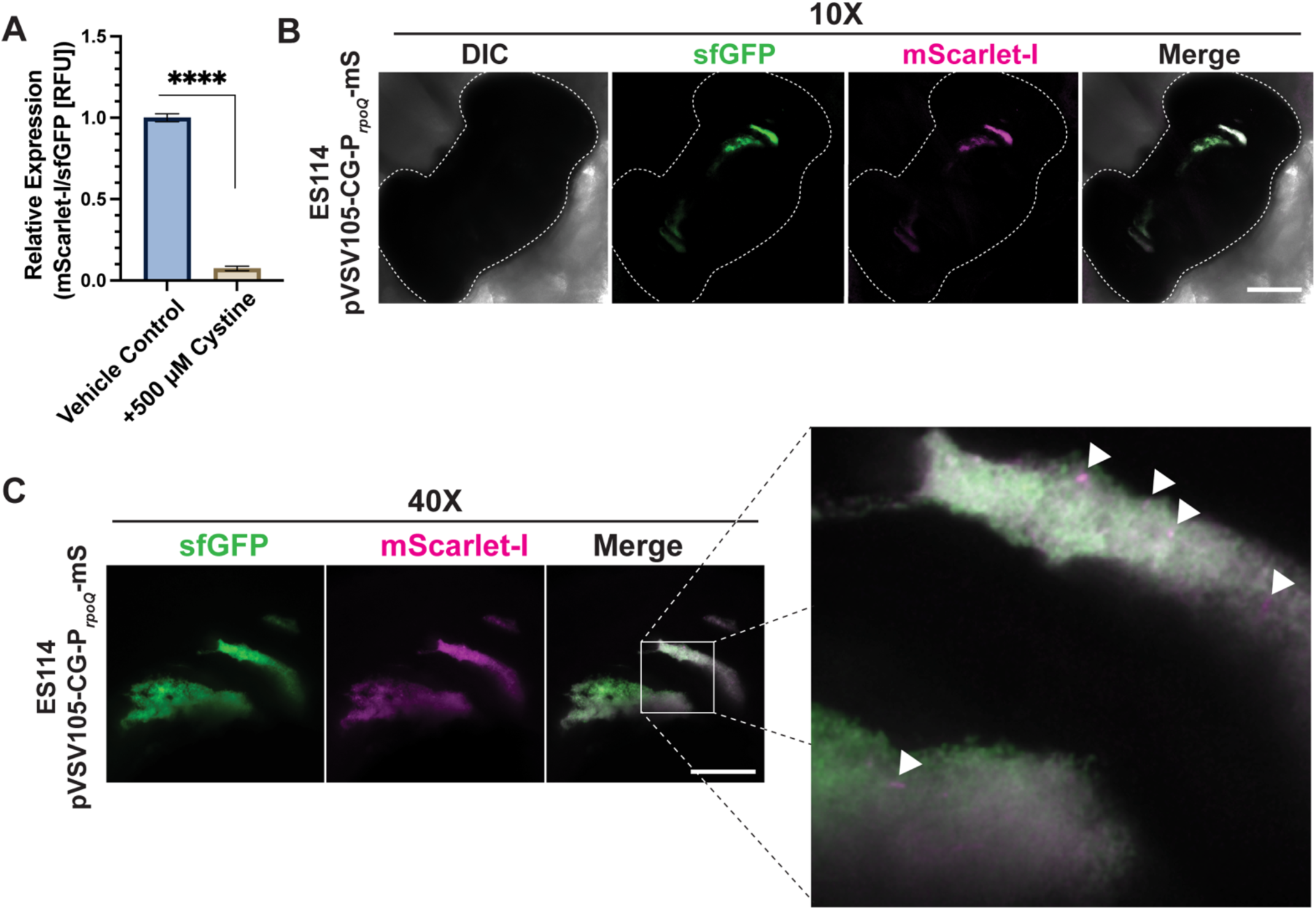
*In vivo* imaging and *in vitro* induction of pVSV105-CG-VmS-derived plasmids in *V. fischeri*. (**A**) *In vitro* repression of pVSV105-CG-P*cysK*-mS by the addition of cystine to the growth media (n = 2 biological replicates). ****, P < 0.0001 by a one-sample *t* test. 10X (**B**) and 40X (**C**) *in vivo* imaging of ES114 pVSV105-CG-P*rpoQ*-mS colonized juvenile *E. scolopes* bobtail squid showing colocalization of sfGFP and mScarlet-I signal in light organ crypt spaces. The light organ and ink sac are outlined with a dotted line in *B*. Scale bars = 150 μm (for B) and 50 μm (for C). Arrows point to areas of concentrated mScarlet-I expression.

Hatchling *E. scolopes* were colonized with *V. fischeri* pVSV105-CG-P*_rpoQ_*-mS for 24 hours prior to fixation and fluorescence microscopy. Robust sfGFP signal was observed within the light organs of colonized hatchings, indicating successful colonization of the reporter strain. In addition, mScarlet-I signal was observed and colocalized with the sfGFP signal, verifying transcriptional activation of the *rpoQ* promoter by *V. fischeri* within the squid light organ (**Fig. 7B**). P*_rpoQ_* activity appeared to be expressed across the entire colonized area, with additional distinct puncta of high reporter signal present, as can be seen in the merged channel image. These pockets of elevated RpoQ activity could be initial areas of colonization, driven by RpoQ gene regulation.

## Discussion

In this study we developed and characterized a dual fluorophore transcriptional and translational reporter vector, pCG-VmS, for use in Gram-negative bacteria, using *P. aeruginosa* and *V. fischeri* as proof-of-concept microorganisms. Utilizing the GO-responsive *arqI* promoter [31]. we established that our vector system can provide robust sensitivity as a transcriptional reporter and allow for visualization of promoter activity in complex environments, such as within a biofilm. We also demonstrated that pCG-VmS could be used to map the cellular localization of proteins without having to add additional dyes to highlight the perimeter of the bacterial cell. The constitutive sfGFP signal allowed for reporter activity normalization, as well as visualization of viable bacteria during microscopy using confocal imaging. Although not shown here, pCG-VmS could potentially be modified to be used with other imaging techniques, such as Stochastic Optical Reconstruction Microscopy (STORM) or Structured Illumination Microscopy (SIM). Also, while we have not constructed these vectors to facilitate bioluminescence or Forster resonance energy transfer experiments (BRET and FRET, respectively), one could easily envision modifying pCG-VmS to accommodate two different protein fusions (sfGFP and mScarlet-I) and use a near-infrared dye to label the bacterial membrane [60]. Alternatively, a third fluorophore could be incorporated into the vector which presents a minimum spectral overlap with the other two signals.

Indeed, a triple reporter system has recently been accomplished to monitor complex mixtures of cyclic nucleotides in *P. aerginosa* cells using mScarlet-I and mGreenLantern as the variable reporter signals and cyan fluorescent protein SCFP3A as a constitutive readout [61]. The fact that this system and the pCG-VmS system described here both utilize mScarlet-I as a readout highlights the superior compatibility of this monomeric, bright fluorophore for use in bacterial imaging.

In this work we also demonstrated that the pCG-VmS fluorescence cassette can be versatile by transferring it to different vector backbones, which enabled its use in different species of Gram-negative bacteria. In our example using *V. fischeri*, transfer of the cassette to a *Vibrio*-specific plasmid allowed for transcriptional reporter activity to be quantified *in vitro* and visualized *in vivo* during colonization of *E. scolopes*. The current plasmid backbone utilized in pCG-VmS, pBBR1, was originally isolated from *Bordetella bronchiseptica* [62], but is usable within a wide array of Gram-negative bacterial genera, including *Escherichia, Rhizobium, Pseudomonas, Salmonella, Klebsiella, Agrobacterium, Azospirullum,* and *Serratia* species [62, 63]. For these and other species, similar to our *V. fisheri* constructions, it is plausible that different constitutive promoters might be required to ‘fine tune’ the normalization signal.

Since the constitutive expression of sfGFP is driven by the *lac*-based promoter *lacUV5*, common knowledge would suggest that robust sfGFP expression would only be possible in species that do not express the *lac* repressor, LacI. Nevertheless, we have observed that despite it possessing an encoded LacI repressor, *E. coli* DH5α harboring pP_A1/04/03_-G-VmS did exhibit a measurable sfGFP fluorescence signal (data not shown). This could be due to the stoichiometry of promoter to LacI within the cell, which would be in favor of the promoter on multi-copy plasmids. Indeed, this effect has been well established in the literature, with an increase in copy number of promoters leading to a decrease in repression efficiency by LacI [64]. Therefore, sufficient sfGFP expression from pP_A1/04/03_-G-VmS may, in fact, be possible even in *lacI*^+^ bacteria. The exact activity of the *lacUV5* promoter in different species can vary, as was the case in *V. fischeri,* which exhibited low sfGFP fluorescence. We attribute this phenomenon to the *E. coli* σ^70^-dependent promoter not being efficiently recognized by the cognate RNAP holoenzyme in this bacterial species. An alternative hypothesis is that LacI homologs from different species might have slightly different DNA recognition motifs, which could affect repression when using the heterologous *lacUV5* promoter sequence [65]. This problem might be circumvented by utilizing the constitutive P_A1/04/03_ promoter, or an option with similar promoter strength, that provides higher expression (as was accomplished in *V. fischeri* here), or simply by changing the LacI recognition site in the *lacUV5* promoter. Taken together, pCG-VmS and pVSV105-CG-VmS have the potential to be useful in many Gram-negative bacterial species but might require fine-tuning expression of sfGFP in accordance with a particular microbe.

A nagging issue with all plasmid-based expression systems is the problem of copy number heterogeneity. A method to lessen the copy number variability of plasmid-based reporters like pCG-VmS would be to construct a reporter that integrates into the chromosome as a single copy. A good system to accomplish this would be the mini-CTX1 system that harbors gentamicin resistance. This system has the ability to integrate as a single-copy into the *P. aeruginosa* genome at the *attB* phage site [66], and could therefore be utilized to make single-copy reporter constructs in *P. aeruginosa*. Indeed, the *attB* site has been previously utilized to constitutively express fluorophores from the P_A1/04/03_ promoter, as well as fluorescent transcriptional reporters [31]. During the construction of pCG-VmS this plasmid, mini-CTX1-Gm-P_A1/04/03_-G-VmS, was created as an intermediate plasmid. However, the feasibility of utilizing mini-CTX1-Gm-P_A1/04/03_-G-VmS in making dual fluorophore reporter constructs *in cis* is currently under investigation and has therefore yet to be confirmed.

Here we also reported the use of pCG-VmS to enable verification of distinct T3SS-off and T3SS-on populations of intracellular *P. aeruginosa* during corneal epithelial cell infections, an observation that had been observed but not verified quantitatively [32, 52]. The study of these distinct populations, especially the T3SS-off bacteria, has been limited by the availability of tools to visualize both bacterial populations concurrently during infection with single-cell accuracy and precise quantitation of the heterogeneous populations. Use of this new tool will allow for further investigation of the underlying mechanisms driving these two distinct populations, as well as possible further dissection of the regulatory mechanisms that control T3SS activation, or repression, during intracellular infection. Moreover, such studies could reveal how the natural spatiotemporal T3SS induction in *P. aeruginosa* during both cellular and animal infections might differ from the more unnatural, *in vitro* induction using EGTA [67].

In summary, the plasmid presented in this study, pCG-VmS, has proven to be a highly useful dual reporter system that expands upon the genetic toolbox available for *P. aeruginosa*. The experiments performed have demonstrated its practicality in a wide variety of applications, including as an *in vitro* and *in vivo* transcriptional reporter and as a translational reporter to determine protein subcellular localization. In addition, pCG-VmS may prove to be an additional tool to study other Gram-negative bacteria. The utility and ease of construction of pCG-VmS-derived reporters should help to further studies of *P. aeruginosa* and other Gram-negative bacteria within the microbiology field.

## Methods

### Ethics Statement

All animal experiments were conducted in accordance with protocols approved by the University of Connecticut Institutional Animal Care and Use Committee (A18-029 and A22-004).

### Bacterial strains and growth conditions

Bacterial strains used in this study can be found in **Table S1***. P. aeruginosa* and *E. coli* cultures were grown in Luria-Bertani media (LB, Difco; Beckton Dickinson, Franklin Lakes, NJ). Liquid cultures were grown at 37°C and 230 rpm on an orbital shaker. *V. fischeri* cultures were grown in LB salt (LBS; 1% tryptone, 0.5% yeast extract, 2% sodium chloride, and 50 mM Tris [pH 7.5]) or Tris-minimal media [68] (TMM, 100 mM Tris pH 7.5, 300 mM NaCl, 50 mM MgSO_4_, 0.33 mM KH_2_PO_4_, 10 μM ferrous ammonium sulfate, 0.1% w/v NH_4_Cl, 10 mM N-acetylglucosamine, 10 mM KCl, and 10 mM CaCl_2_) at 28°C. Culture density was monitored using a Genesys 150 UV-Vis spectrophotometer (Thermo Fisher Scientific, Waltham, MA) at a wavelength of 600 nm. Solid media was solidified with 1.5% agar (bacteriological; VWR, Solon, OH). Super optimal broth with catabolite repression (SOC) was made with 20 g/L tryptone, 5 g/L yeast extract, 0.5 g/L NaCl, 10 mM MgCl_2_, 10 mM MgSO_4_, 2.5 mM KCl, and 10 mM glucose. Antibiotic concentrations for *E. coli* were as follows: 15 μg/mL Gentamicin (Gm), 15 μg/mL chloramphenicol (Cm). For *P. aeruginosa,* Gm was used at 30 μg/mL. For *V. fischeri,* Cm was used at a concentration of 5 μg/mL.

### Fluorescence reporter assays

For *P. aeruginosa* fluorescence reporter assays, experiments were carried out as previously described [31]. Briefly, overnights of *P. aeruginosa* MPAO1 carrying the indicated plasmids were subcultured 1:100 from overnights into fresh LB media. Cultures were then grown at 37°C and 230 rpm for 2 hours before addition of compounds. Cultures were grown for an additional 3 hours before fluorescence readings were taken using a BioTek H1 Synergy multimode plate reader (BioTek, Winooski, VT). Green (excitation 485/20 nm, emission 528/20 nm, using a dichroic mirror 510 nm) and Red (excitation 575/15 nm, emission 635/32 nm, using a dichroic mirror 595 nm) filter cubes were used for fluorescence readings. For *V. fischeri* reporter constructs, experiments were performed as in ref. [57] with minor modifications. Briefly, *V. fischeri* strains were inoculated from freezer stocks into TMM media and grown at 28°C for several hours. Strains were then subcultured 1:100 into fresh TMM supplemented with HCl vehicle or 500 μM cystine and grown overnight. Cultures were pelleted and resuspended in PBS before fluorescence readings were performed as per *P. aeruginosa* protocol stated above.

### P. aeruginosa growth assays

Growth assays were done as described previously [31, 33]. Briefly, 1 mL cultures were washed one time in fresh LB media. Cultures were then normalized to an OD_600_ of 0.05 in fresh media. 200 μL of cultures were then aliquoted in triplicate into blacked-walled 96 well plates (655096; Greiner Bio-One, Stonehouse, UK). Growth and fluorescence were then monitored over time in a BioTek H1 Synergy multimode plate reader (BioTek, Winooski, VT) at 37°C with 567 cpm linear shaking.

### Live cell fluorescence microscopy for protein subcellular localization

Microscopy was performed as described previously [31]. Briefly, an overnight culture of *P. aeruginosa* MPAO1 containing the pCG-VmS derived reporter plasmid was subcultured 1:100 into fresh medium and grown for 2 hours at 37°C and 230 rpm. The culture was diluted 1:2 in fresh LB medium and 1-2 μL of cells were then transferred to a 1.5% agarose LB (LBA) pad on a cleaned slide. LBA Pads were made by pipetting 50 μL of 1.5% agarose LB into a 125 μL Gene Frame (AB0578; Thermo Scientific) and placing a clean slide on top until the media solidified. The pad was then sealed with a No. 1.5 coverslip before imaging.

### Construction of pCG-VmS

To construct pCG-VmS, mini-CTX1-Gm-mCherry was digested with NcoI and KpnI to remove the copy of mCherry. A linear DNA fragment synthesized by IDT containing the mScarlet-I coding sequence was PCR amplified with primer pairs NcoI-mScarlet-I-F1/KpnI-mScarlet-I-R1 and NcoI-mScarlet-I-F2/KpnI-mScarlet-I-R2 (**Table S1**) and ligated into the digested plasmid to create miniCTX1-Gm-mScarlet-I. To insert the _PA1/04/03_-sfGFP cassette and 150 bp of randomly selected DNA sequence to space the divergent sfGFP and mScarlet-I genes, the reverse complement of the P_A1/04/03_-sfGFP was amplified from mini-CTX1-Gm-P_A1/04/03_-sfGFP with a 3’ sequence complementary to a noncoding section of DNA from pUC18T-mini-Tn7T-Gm with primers SacI-sfGFP-revcomp-F1 and P_A1/04/03_-150bp-pUC18T-R. 150 bp of random DNA sequence from pUC18T-mini-Tn7T-Gm was amplified using primer pairs P_A1/04/03_-150bp-pUC18T-F and NotI-150bp-pUC18T-R1, followed by splicing by overlap extension with the previously mentioned PCR product. The resulting gel-extracted DNA product was reamplified with primer pairs SacI-sfGFP-revcomp-F1/NotI-150bp-pUC18T-R2 and SacI-sfGFP-revcomp-F2/NotI-150bp-pUC18T-R1 and ligated into SacI and NotI digested miniCTX1-Gm-mScarlet-I to create mini-CTX1-PA_1/04/03_-G-mS.

The PA_1/04/03_-sfGFP-MCS-mScarlet-I cassette was amplified using primer pairs SacI-sfGFP-revcomp-F1/KpnI-mScarlet-I-R2 and SacI-sfGFP-revcomp-F2/KpnI-mScarlet-I-R1 and inserted into SacI and KpnI digested pCC21 to create pP_A1/04/03_-G-VmS. The P_A1/04/03_ promoter was then swapped for the lacUV5 promoter by digesting pP_A1/04/03_-G-VmS with PfoI and SapI and insertion of SapI and PfoI digested P_lacUV5_ gene fragment (synthesized by IDT) to create pP_lacUV5_-G-VmS. Finally, an rrnB terminator was amplified using primer pairs rrnBTerm-Gibson-F/rrnBTerm-Gibson-R from pMMB67EH and inserted at the PfoI site between the divergent genes in the direction of the sfGFP coding sequence by Gibson assembly to yield pCG-VmS.

Transcriptional fusions were made in pCG-VmS by amplification of the gene of interest upstream regulatory region with primers containing 5’ BamHI and 3’ HindIII overhangs and insertion into BamHI and HindIII digested pCG-VmS. Primer pairs for P*_arqI_* (145 bp): BamHI-P_ArqI_-F1/HindIII-P_ArqI_-R2 and BamHI-P_arqI_-F2/HindIII-P_ArqI_-R1. Primer pairs for P_exoS_ amplification from pJNE05 [69]: BamHI-P_exoS_-F1/HindIII-P_exoS_-R2 and BamHI-P_exoS_-F2/HindIII-P_exoS_-R1. Translational fusions of ArqI to mScarlet-I were made by amplification of the *arqI* upstream regulatory region (145 bp), *arqI* gene, and a C-terminal domain breaking linker [27] from pSB109-ArqI-sfGFP using primer pairs BamHI-P_arqI_-F1/NcoI-Linker-R2 and BamHI-P_ArqI_-F1/NcoI-LinkerR1. The resulting product was ligated into BamHI and NcoI digested pCG-VmS to yield pCG-P*_arqI_*-ArqI-mS.

### Construction of flow cytometry control plasmids

For flow cytometry controls, pSB109-sfGFP and pSB109-mScarlet-I were constructed. The sfGFP coding sequence was amplified using primer pairs NcoI-sfGFP-F1/NdeI-sfGFP-R2 and NcoI-sfGFP-F2/NdeI-sfGFP-R1. The resulting product was then ligated into NcoI/NdeI digested pSB109 using a restrictionless cloning method [70]. Similarly, the mScarlet-I coding sequence was amplified with primer pairs NcoI-mScarlet-I-F1/NdeI-mScarlet-I-R2 and NcoI-mScarlet-I-F2/NdeI-mScarlet-I-R1 and ligated into pSB109.

### Construction of pVSV105-CG-VmS

To construct pVSV105-CG-VmS, the native NcoI site in pVSV105 was disrupted using site directed mutagenesis by the Quick Change Method [71] using primers pVSV105-NcoI-Poison-F and pVSV105-NcoI-Poison-R (**Table S1**). The resulting plasmid was digested by NotI and SphI followed by insertion of a PCR amplified copy of P_A1/04/03_-sfGFP from pP_A1/04/03_-G-VmS using primer pairs NotI-sfGFP-F1/SphI-sfGFP-R2 and NotI-sfGFP-F2/SphI-sfGFP-R1 to yield pVSV105-CG. pVSV105-CG was digested with SacI and SpeI before insertion of a PCR amplified copy of the RBS and mScarlet-I genes amplified from pP_A1/04/03_-G-VmS using primer pairs SacI-RBS-mScarlet-I-F1/SpeI-mScarlet-R2 and SacI-RBS-mScarlet-I-F2/SpeI-mScarlet-R1 to yield pVSV105-CG-VmS. Transcriptional fusions were made by insertion of upstream regulatory regions at the SphI and KpnI sites using the following primers. P_rpoQ_: SphI-P_rpoQ_-F/KpnI-P_rpoQ_-R, P_cysK_: SphI-P_cysK_-F/KpnI-P_cysK_-R (**Table S1**).

### Flow Cytometry

Cultures were grown as described in the ‘Fluorescence reporter assays’ section. Samples were diluted to a cell density of ∼5 x 10^7^ CFU/mL (OD_600_ 0.1 = 1 x 10^8^ CFU/mL) in 2 mL PBS. Samples were read on a full-spectrum cytometer, the 5-laser Aurora (Cytek, Fremont, CA). Appropriate sfGFP and mScarlet-I single color controls were used to unmix and gate the data. Analyses of the unmixed data were performed in FlowJo v10.9.0 (BD Life Sciences, Ashland, OR). sfGFP-positive cells were gated on for all analyses.

### Biofilm flow cell imaging

Flow cell bioreactors were prepared as previously described [72]. Log-phase MPAO1 pCG-ParqI-mS cultures grown in tryptic soy broth (TSB) were diluted to 0.01 OD_600_ in 1% TSB. Diluted cultures were injected into continuous flow cell chambers and incubated and then inverted for 1 hour at room temperature to allow for cell attachment. Following attachment, continuous flow of 1% TSB media was supplied at a rate of 10 mL/h for 96 hours at room temperature. Resultant biofilms were treated with 10 mM GO in 1% TSB under 10 mL/h flow for 3 hours and subsequently imaged using confocal microscopy (Nikon A1R). Three images were taken for each sample in 3 independent experiments. Representative z-stack images were processed in Volocity Image Analysis (Improvision, Coventry, UK).

### Squid Colonization Experiments

Hatchling *E. scolopes* were colonized with *V. fischeri* KV10353 by addition of 3,000 colony forming units per milliliter (CFUs/ml) to filter-sterilized artificial seawater (FSASW) for 3 hours followed by transfer to 20 ml glass scintillation vials containing FSASW. Light organ colonization was determined at 24 hours post inoculation by monitoring luminescence using a Turner TD 202/20 single tube luminometer (Turner Instruments). Aposymbiotic control animals were maintained in FSASW without the addition of *V. fischeri* KV10353. Symbiotic animals were anaesthetized in 2% ethanol in FSAWSW and then fixed for 12 hours in 4% paraformaldehyde in FSASW at 4°C. Fixed animals were then rinsed and stored in FSASW until imaging.

For imaging, animals were mounted in 1X marine phosphate buffered saline (50 mM sodium phosphate, 0.45 M NaCl, pH 7.4) on concavity well slides (Electron Microscopy Sciences). Mounted animals were then imaged using a Nikon A1R laser-scanning confocal microscope with 10X 0.30 NA plan fluorite and 40X 1.20 NA plan apochromat objectives at the University of Connecticut Advanced Light Microscopy Facility. Images for sfGFP and mScarlet-I signals were acquired using 488-and 561-nm laser lines, respectively, to excite the fluorophores.

### Cell culture and Time Lapse Imaging

hTCEpi cells were maintained in KGM-2 media (PromoCell) [73]. HeLa cells were maintained in phenol red-free DMEM (Gibco) containing 10% FBS. One day preceding experiments, hTCEpi cells were plated on No. 1.5 glass-bottom 24 well plates (MatTek) in KGM-2 at 75% confluence with 1.06 mM calcium to induce differentiation [73]. HeLa cells were plated at 50% confluence on optical plastic 8-well chambered coverslips (Ibidi). For infection, bacterial suspensions were made in PBS from 16-hour lawns grown on TSA media containing 100 µg/ml gentamicin at 37°C. A multiplicity of infection equal to 10 was calculated using OD_540_ of 1 equal to 4 x 10^8^ CFU per mL. Bacteria were added directly to culture media and allowed to invade cells. Media was replaced with media containing amikacin (200 µg/ml), or additional polymyxin B (10 µg/ml) at 3 hours post infection to kill extracellular bacteria. Of note, this timepoint precedes observed fluorescence of reporters for ExoS, and this technique has been validated to observe intracellular bacteria previously [41]. Beginning at 3.5 hours post infection, images were captured on a Ti2-E inverted microscope with X-Cite XYLIS XT720S Broad Spectrum LED Illumination System equipped with an Okolab stage-top incubation chamber to maintain 37°C and 5% CO_2_, a DS-Qi2 CMOS camera, and CFI Plan Apochromat Lambda D 40X air NA 0.95 objective. Time lapse fields were selected without observing fluorescence channels to limit bias in field selection. Images were captured with intervals of 5 or 10 minutes.

## Acknowledgements

This work was funded though National Institute of Health (NIH) grant R01 GM141230 and NIAID R01 A135060 to A.T.U., grant NIH R01 EY034239 to A.R.K., grant P20 GM103440 and R16 GM149513 to B.S.T., grant R35 GM130355 to K.L.V., and National Science Foundation (NSF) grant IOS-IOS-2247195 and Gordon and Betty Moore Foundation grants 9349 and 12342 to S.V.N.

## Author contributions

C.J.C. and D.G.G conceived the project, C.J.C., D.G.G., Z.J.R., E.K.C., D.L.K., S.V.N., B.S.T. performed the experiments, C.J.C., D.G.G., Z.J.R., E.K.C., D.L.K., S.V.N., B.S.T., A.R.K., K.L.V., A.T.U. analyzed the data and interpreted the results, and C.J.C. and A.T.U. wrote the manuscript.

## Resource availability

### Lead contact

Further information and requests for resources and reagents should be directed to and will be fulfilled by the lead contact, Andrew T. Ulijasz

### Materials availability

All reagents are available by contacting the lead contact of this study. Unique reagents by MTA.

### Declaration of generative AI and AI-assisted technologies

No AI was used in the writing or data involved in this project.

## References

1. Rodriguez, E.A., et al., The Growing and Glowing Toolbox of Fluorescent and Photoactive Proteins. Trends in Biochemical Sciences, 2017. 42(2): p. 111–129.

2. Chudakov, D.M., et al., Fluorescent Proteins and Their Applications in Imaging Living Cells and Tissues. Physiological reviews., 2010. 90(3): p. 1103–1163.

3. He, L., et al., In vivo study of gene expression with an enhanced dual-color fluorescent transcriptional timer. ELife., 2019. 8.

4. Ceolin, S., et al., A sensitive mNeonGreen reporter system to measure transcriptional dynamics in Drosophila development. Communications Biology, 2020. 3(1).

5. Tanz, S.K., et al., Fluorescent protein tagging as a tool to define the subcellular distribution of proteins in plants. Frontiers in plant science., 2013. 4.

6. Stadler, C., et al., Immunofluorescence and fluorescent-protein tagging show high correlation for protein localization in mammalian cells. Nature methods., 2013. 10(4): p. 315–323.

7. Speare, L., et al., Bacterial symbionts use a type VI secretion system to eliminate competitors in their natural host. Proceedings of the National Academy of Sciences, 2018. 115(36): p. E8528–E8537.

8. Schlechter, R.O., et al., Fluorescent Protein Expression as a Proxy for Bacterial Fitness in a High-Throughput Assay. Applied and environmental microbiology : AEM., 2021. 87(18).

9. Sekar, R.B. and A. Periasamy, Fluorescence resonance energy transfer (FRET) microscopy imaging of live cell protein localizations. The Journal of Cell Biology, 2003. 160(5): p. 629–633.

10. Fang, C., Y. Huang, and Y. Zhao, Review of FRET biosensing and its application in biomolecular detection. American journal of translational research., 2023. 15(2): p. 694–709.

11. Zhang, Y., et al., Fast and sensitive GCaMP calcium indicators for imaging neural populations. Nature, 2023. 615(7954): p. 884–891.

12. Glanville, D.G., et al., A high-throughput method for identifying novel genes that influence metabolic pathways reveals new iron and heme regulation in Pseudomonas aeruginosa. mSystems, 2021. 6(1).

13. Stream, A. and C.A. Madigan, Zebrafish: an underutilized tool for discovery in host-microbe interactions. Trends Immunol, 2022. 43(6): p. 426–437.

14. Jiang, M., et al., A Red Fluorescent Protein Reporter System Developed for Measuring Gene Expression in Photosynthetic Bacteria under Anaerobic Conditions. Microorganisms, 2022. 10(2): p. 201.

15. Uliczka, F., et al., Monitoring of Gene Expression in Bacteria during Infections Using an Adaptable Set of Bioluminescent, Fluorescent and Colorigenic Fusion Vectors. PLoS ONE, 2011. 6(6): p. e20425.

16. Singh, K.P., et al., Mechanisms and Measurement of Changes in Gene Expression. Biological Research For Nursing, 2018. 20(4): p. 369–382.

17. Wilson, E., et al., Using Fluorescence Intensity of Enhanced Green Fluorescent Protein to Quantify Pseudomonas aeruginosa. Chemosensors, 2018. 6(2): p. 21.

18. Million-Weaver, S., et al., Quantifying Plasmid Copy Number to Investigate Plasmid Dosage Effects Associated with Directed Protein Evolution, in Methods in Molecular Biology. 2012, Springer New York. p. 33–48.

19. Vrla, G.D., et al., Cytotoxic alkyl-quinolones mediate surface-induced virulence in Pseudomonas aeruginosa. PLOS Pathogens, 2020. 16(9): p. e1008867.

20. Yu, C.-S., C.-J. Lin, and J.-K. Hwang, Predicting subcellular localization of proteins for Gram-negative bacteria by support vector machines based on n-peptide compositions. Protein Science, 2004. 13(5): p. 1402–1406.

21. Rudner, D.Z. and R. Losick, Protein Subcellular Localization in Bacteria. Cold Spring Harbor Perspectives in Biology, 2010. 2(4): p. a000307–a000307.

22. Pi, H., et al., Clostridioides difficile ferrosome organelles combat nutritional immunity. Nature., 2023. 623(7989): p. 1009–1016.

23. Yoon, S.A., et al., Strategies of Detecting Bacteria Using Fluorescence-Based Dyes. Frontiers in chemistry., 2021. 9.

24. Heinrich, K., et al., Molecular Basis and Ecological Relevance of Caulobacter Cell Filamentation in Freshwater Habitats. mBio, 2019. 10(4).

25. Banks, E.J., et al., Asymmetric peptidoglycan editing generates cell curvature in Bdellovibrio predatory bacteria. Nature Communications, 2022. 13(1).

26. Obara, B., et al., Bacterial cell identification in differential interference contrast microscopy images. BMC Bioinformatics, 2013. 14(1): p. 134.

27. Kjos, M., et al., Bright Fluorescent Streptococcus pneumoniae for Live-Cell Imaging of Host-Pathogen Interactions. Journal of Bacteriology, 2015. 197(5): p. 807–818.

28. Berlec, A., et al., In vivo imaging of Lactococcus lactis, Lactobacillus plantarum and Escherichia coli expressing infrared fluorescent protein in mice. Microb Cell Fact, 2015. 14: p. 181.

29. Hall, C., et al., iRFP (near-infrared fluorescent protein) imaging of subcutaneous and deep tissue tumours in mice highlights differences between imaging platforms. Cancer Cell Int, 2021. 21(1): p. 247.

30. Barbier, M. and F.H. Damron, Rainbow Vectors for Broad-Range Bacterial Fluorescence Labeling. PLoS One, 2016. 11(3): p. e0146827.

31. Corcoran, C.J., et al., A glyoxal-specific aldehyde signaling axis in Pseudomonas aeruginosa that influences quorum sensing and infection. Nat Commun, 2025. 16(1): p. 6616.

32. Kumar, N.G., et al., Pseudomonas aeruginosa Can Diversify after Host Cell Invasion to Establish Multiple Intracellular Niches. MBio., 2022. 13(6).

33. Corcoran, C.J., et al., The glyoxal response in Pseudomonas aeruginosa represses PQS-mediated communication through a novel regulatory axis. Unpublished, 2023.

34. Jahn, M., et al., Copy number variability of expression plasmids determined by cell sorting and Droplet Digital PCR. Microbial Cell Factories, 2016. 15(1).

35. Bindels, D.S., et al., mScarlet: a bright monomeric red fluorescent protein for cellular imaging. Nature methods., 2017. 14(1): p. 53–56.

36. Balleza, E., J.M. Kim, and P. Cluzel, Systematic characterization of maturation time of fluorescent proteins in living cells. Nature Methods, 2018. 15(1): p. 47–51.

37. Lanzer, M. and H. Bujard, Promoters largely determine the efficiency of repressor action. Proceedings of the National Academy of Sciences of the United States of America., 1988. 85(23): p. 8973–8977.

38. Lambertsen, L., C. Sternberg, and S. Molin, Mini-Tn7 transposons for site-specific tagging of bacteria with fluorescent proteins. Environmental Microbiology, 2004. 6(7): p. 726–732.

39. Wang, B., et al., Pseudomonas aeruginosa PA14 produces R-bodies, extendable protein polymers with roles in host colonization and virulence. Nature Communications, 2021. 12(1).

40. Klausen, M., et al., Biofilm formation by Pseudomonas aeruginosa wild type, flagella and type IV pili mutants. Molecular Microbiology, 2003. 48(6): p. 1511–1524.

41. Kroken, A.R., et al., The Impact of ExoS on Pseudomonas aeruginosa Internalization by Epithelial Cells Is Independent of fleQ and Correlates with Bistability of Type Three Secretion System Gene Expression. mBio, 2018. 9(3).

42. Orosz, A., I. Boros, and P. Venetianer, Analysis of the complex transcription termination region of the Escherichia coli rrnB gene. European Journal of Biochemistry, 1991. 201(3): p. 653–659.

43. Shcherbakova, D.M., O.M. Subach, and V.V. Verkhusha, Red Fluorescent Proteins: Advanced Imaging Applications and Future Design. Angewandte Chemie International Edition, 2012. 51(43): p. 10724–10738.

44. Helaine, S., et al., Dynamics of intracellular bacterial replication at the single cell level. Proceedings of the National Academy of Sciences, 2010. 107(8): p. 3746–3751.

45. Fleiszig, S.M., T.S. Zaidi, and G.B. Pier, Pseudomonas aeruginosa invasion of and multiplication within corneal epithelial cells in vitro. Infect Immun, 1995. 63(10): p. 4072–7.

46. Angus, A.A., et al., Pseudomonas aeruginosa induces membrane blebs in epithelial cells, which are utilized as a niche for intracellular replication and motility. Infect Immun, 2008. 76(5): p. 1992–2001.

47. Kroken, A.R., et al., Exotoxin S secreted by internalized Pseudomonas aeruginosa delays lytic host cell death. PLoS Pathog, 2022. 18(2): p. e1010306.

48. Kumar, N.G., et al., Pseudomonas aeruginosa Can Diversify after Host Cell Invasion to Establish Multiple Intracellular Niches. mBio, 2022. 13(6): p. e0274222.

49. Urbanowski, M.L., E.D. Brutinel, and T.L. Yahr, Translocation of ExsE into Chinese hamster ovary cells is required for transcriptional induction of the Pseudomonas aeruginosa type III secretion system. Infect Immun, 2007. 75(9): p. 4432–9.

50. Kroken, A.R., et al., Intracellular replication of Pseudomonas aeruginosa in epithelial cells requires suppression of the caspase-4 inflammasome. mSphere, 2023: p. e0035123.

51. McMackin, E.A.W., et al., Cautionary Notes on the Use of Arabinose-and Rhamnose-Inducible Expression Vectors in Pseudomonas aeruginosa. J Bacteriol, 2021. 203(16): p. e0022421.

52. Resko, Z.J., et al., Evidence for intracellular Pseudomonas aeruginosa. J Bacteriol, 2024. 206(5): p. e0010924.

53. Visick, K.L., E.V. Stabb, and E.G. Ruby, A lasting symbiosis: how Vibrio fischeri finds a squid partner and persists within its natural host. Nature reviews., 2021. 19(10): p. 654–665.

54. Nyholm, S.V. and M.J. Mcfall-Ngai, A lasting symbiosis: how the Hawaiian bobtail squid finds and keeps its bioluminescent bacterial partner. Nature Reviews Microbiology, 2021. 19(10): p. 666–679.

55. Dunn, A.K., et al., New rfp-and pES213-derived tools for analyzing symbiotic Vibrio fischeri reveal patterns of infection and lux expression in situ. Applied and Environmental Microbiology, 2006. 72(1): p. 802–810.

56. Ludvik, D.A., et al., Hybrid Histidine Kinase BinK Represses Vibrio fischeri Biofilm Signaling at Multiple Developmental Stages. Journal of bacteriology : JB., 2021. 203(15).

57. Wasilko, N.P., et al., Sulfur availability for Vibrio fischeri growth during symbiosis establishment depends on biogeography within the squid light organ. Molecular Microbiology, 2019. 111(3): p. 621–636.

58. Cao, X., et al., The Novel Sigma Factor-Like Regulator RpoQ Controls Luminescence, Chitinase Activity, and Motility in Vibrio fischeri. MBio., 2012. 3(1): p. e00285.

59. Lupp, C. and E.G. Ruby, Vibrio fischeri Uses Two Quorum-Sensing Systems for the Regulation of Early and Late Colonization Factors. Journal of Bacteriology, 2005. 187(11): p. 3620–3629.

60. Lucidi, M., et al., Expanding the microbiologist toolbox via new far-red-emitting dyes suitable for bacterial imaging. Microbiol Spectr, 2024. 12(1): p. e0369023.

61. Zheng, X., et al., The surface interface and swimming motility influence surface-sensing responses in Pseudomonas aeruginosa. Proc Natl Acad Sci U S A, 2024. 121(39): p. e2411981121.

62. Antoine, R. and C. Locht, Isolation and molecular characterization of a novel broad-host-range plasmid from Bordetella bronchiseptica with sequence similarities to plasmids from Gram-positive organisms. Molecular Microbiology, 1992. 6(13): p. 1785–1799.

63. Szpirer, C.D.Y., M. Faelen, and M. Couturier, Mobilization Function of the pBHR1 Plasmid, a Derivative of the Broad-Host-Range Plasmid pBBR1. Journal of Bacteriology, 2001. 183(6): p. 2101–2110.

64. Glascock, C.B. and M. J Weickert, Using chromosomal lacIQ1 to control expression of genes on high-copy-number plasmids in Escherichia coli1Published in conjunction with A Wisconsin Gathering Honoring Waclaw Szybalski on the occasion of his 75th year and 20 years of Editorship-in-Chief of Gene, 10–11 August 1997, University of Wisconsin, Madison, WI, USA.1. Gene., 1998. 223(1-2): p. 221–231.

65. Falcon, C.M. and K.S. Matthews, Operator DNA sequence variation enhances high affinity binding by hinge helix mutants of lactose repressor protein. Biochemistry, 2000. 39(36): p. 11074–83.

66. Hoang, T.T., et al., Integration-Proficient Plasmids for Pseudomonas aeruginosa: Site-Specific Integration and Use for Engineering of Reporter and Expression Strains. Plasmid : a journal of molecular genetics with emphasis on plasmid biology., 2000. 43(1): p. 59–72.

67. Dasgupta, N., et al., Transcriptional induction of the Pseudomonas aeruginosa type III secretion system by low Ca2+ and host cell contact proceeds through two distinct signaling pathways. Infect Immun, 2006. 74(6): p. 3334–41.

68. Christensen, D.G. and K.L. Visick, Vibrio fischeri : Laboratory Cultivation, Storage, and Common Phenotypic Assays. Current protocols in microbiology., 2020. 57(1).

69. Brutinel, E.D., et al., Characterization of ExsA and of ExsA-dependent promoters required for expression of the Pseudomonas aeruginosa type III secretion system. Molecular microbiology., 2008. 68(3): p. 657–671.

70. Ulijasz, A.T., A. Grenader, and B. Weisblum, A vancomycin-inducible lacZ reporter system in Bacillus subtilis: induction by antibiotics that inhibit cell wall synthesis and by lysozyme. Journal of Bacteriology, 1996. 178(21): p. 6305–6309.

71. Liu, H. and J.H. Naismith, An efficient one-step site-directed deletion, insertion, single and multiple-site plasmid mutagenesis protocol. BMC Biotechnology, 2008. 8(1): p. 91.

72. Christensen, B.B., et al., [2] Molecular tools for study of biofilm physiology. Biofilms, 1999. 310: p. 20–42.

73. Robertson, D.M., et al., Characterization of growth and differentiation in a telomerase-immortalized human corneal epithelial cell line. Invest Ophthalmol Vis Sci, 2005. 46(2): p. 470–8.

