## Supplemental Figure Legends for "A versatile dual-color bacterial reporter system highlights two distinct *Pseudomonas aeruginosa* Type 3 secretion system intracellular populations"

**Supplemental Movie 1. Optimization of a constitutive GFP promotor**. PAO1F∆*exoSTY* was transformed with either PA_1/04/03_ (labeled “version 1”) or pCG-VmS (labeled “version 2”), each encoding the promotor for ExoS upstream of mScarlet-I, later termed pCG-P*_exoS_*-mS. Bacteria were incubated with HeLa cells for 3 hours to allow internalization. Planktonic bacteria were removed and media replaced with amikacin-containing media. Time-lapse images were captured every 10 minutes to track intracellular bacterial replication fluorophore expression. GFP and mScarlet-I signal were scaled identically in these visualizations.

**Supplemental Movie 2. Comparison of pCG-P*_exoS_*-mS to a previously characterized reporter for T3SS activity.** PAO1F was transformed with either PJNE05 (encodes the promotor for ExoS upstream of GFP) or pCG-P*_exoS_*-mS. Bacteria were incubated with hTCEpi cells for 3 hours to allow internalization. Planktonic bacteria were removed and media replaced with amikacin-containing media. Time-lapse images were captured every 5 minutes to compare the two fluorescent reporters in intracellular populations.

**Supplemental Movie 3. Amikacin and Polymyxin B eliminate extracellular bacteria.** PAO1F transformed with pCG-P*_exoS_*-mS was incubated with hTCEpi cells for 3 hours to allow internalization. Planktonic bacteria were removed and media replaced with amikacin and polymyxin B-containing media, which causes extracellular bacteria to lyse and diminishes their fluorescence within 1-2 hours. Time-lapse images were captured every 5 minutes.
